# Movements of *Mycoplasma mobile* gliding machinery detected by high-speed atomic force microscopy

**DOI:** 10.1101/2021.01.28.428740

**Authors:** Kohei Kobayashi, Noriyuki Kodera, Taishi Kasai, Yuhei O Tahara, Takuma Toyonaga, Masaki Mizutani, Ikuko Fujiwara, Toshio Ando, Makoto Miyata

## Abstract

*Mycoplasma mobile*, a parasitic bacterium, glides on solid surfaces, such as animal cells and glass by a special mechanism. This process is driven by the force generated through ATP hydrolysis on an internal structure. However, the spatial and temporal behaviors of the internal structures in living cells are unclear. In this study, we detected the movements of the internal structure by scanning cells immobilized on a glass substrate using high-speed atomic force microscopy (HS-AFM). By scanning the surface of a cell, we succeeded in visualizing particles, 2 nm in hight and aligned mostly along the cell axis with a pitch of 31.5 nm, consistent with previously reported features based on electron microscopy. Movements of individual particles were then analyzed by HS-AFM. In the presence of sodium azide, the average speed of particle movements was reduced, suggesting that movement is linked to ATP hydrolysis. Partial inhibition of the reaction by sodium azide enabled us to analyze particle behavior in detail, showing that the particles move 9 nm right, relative to the gliding direction, and 2 nm into the cell interior in 330 ms, then return to their original position, based on ATP hydrolysis.

**IMPORTANCE:** The *Mycoplasma* genus contains bacteria generally parasitic to animals and plants. Some *Mycoplasma* species form a protrusion at a pole, bind to solid surfaces, and glide by a special mechanism linked to their infection and survival. The special machinery for gliding can be divided into surface and internal structures that have evolved from rotary motors represented by ATP synthases. This study succeeded in visualizing the real-time movements of the internal structure by scanning from the outside of the cell using an innovative high-speed atomic force microscope, and then analyzing their behaviors.

## INTRODUCTION

Many bacteria translocate to nutrient-rich places and escape from repellent substances by manipulating external appendages, such as flagella and pili (1, 2). However, class *Mollicutes*, a small group of bacteria, have as many as three of their own motility mechanisms. Class Mollicutes evolved from phylum *Firmicutes* by losing peptidoglycan synthesis and flagella swimming to evade host innate immunity in their parasitic life (1). Among *Mollicutes*, the gliding motility of *Mycoplasma mobile*, the subject in this study, is suggested to have evolved from a combination of ATP synthase and cell adhesion (1, 3-8).

*M. mobile*, isolated from a freshwater fish, is a flask-shaped bacterium with a length of 0.8 μm (Fig. 1A). *M. mobile* glides in the direction of tapered end on solid surfaces, such as animal cells, glass, and plastics. Its gliding speed is 2.5-4 μm/s, which is 3-7 times its own cell length (6, 9). The gliding machinery is divided into surface and internal structures, both of which are composed of 450 units (Fig. 1A) (3, 4, 6, 10). The internal structure is characterized by multiple chains. An *M. mobile* cell has approximately 28 chains around the base of protrusion (Fig. 1A). Each chain consists of uniformly-sized particles, which are 13 nm in width and 21 nm in length (4). Interestingly, the amino acid sequence of component proteins suggests that this chain structure has evolved from ATP synthase (3, 4, 6, 8, 11). Recently, the isolated internal structure was shown to hydrolyze ATP through conformational changes, suggesting that the internal structure functions as a motor and generates the force for gliding (4, 6). The surface structure is composed of three large proteins, Gli349, Gli521, and Gli123. Gli349 has a binding site for sialylated oligosaccharide at its tip and plays the role of a “leg” in gliding (5, 12-16). Gli521 and Gli123 have been proposed to act as a “crank” that transmits force (17-20) and as a “mount” to correctly localize the surface proteins (15). A working model for the gliding mechanism has been suggested as follows (4, 6, 9, 21): the force for gliding generated based on ATP-derived energy by the special motor is transmitted across the membrane to the surface structure, including the leg structure. Then, the foot (the tip structure of the leg) repeatedly catches, pulls, and releases the sialylated oligosaccharides (5, 12), the major structures on host animal surfaces (22-24), resulting in cell migration (17, 25-28). This explains the gliding mechanism at the bacterial surface; however, the spatial and temporal behaviors and movements of internal motors in living cells have not been examined.

**FIG 1.**
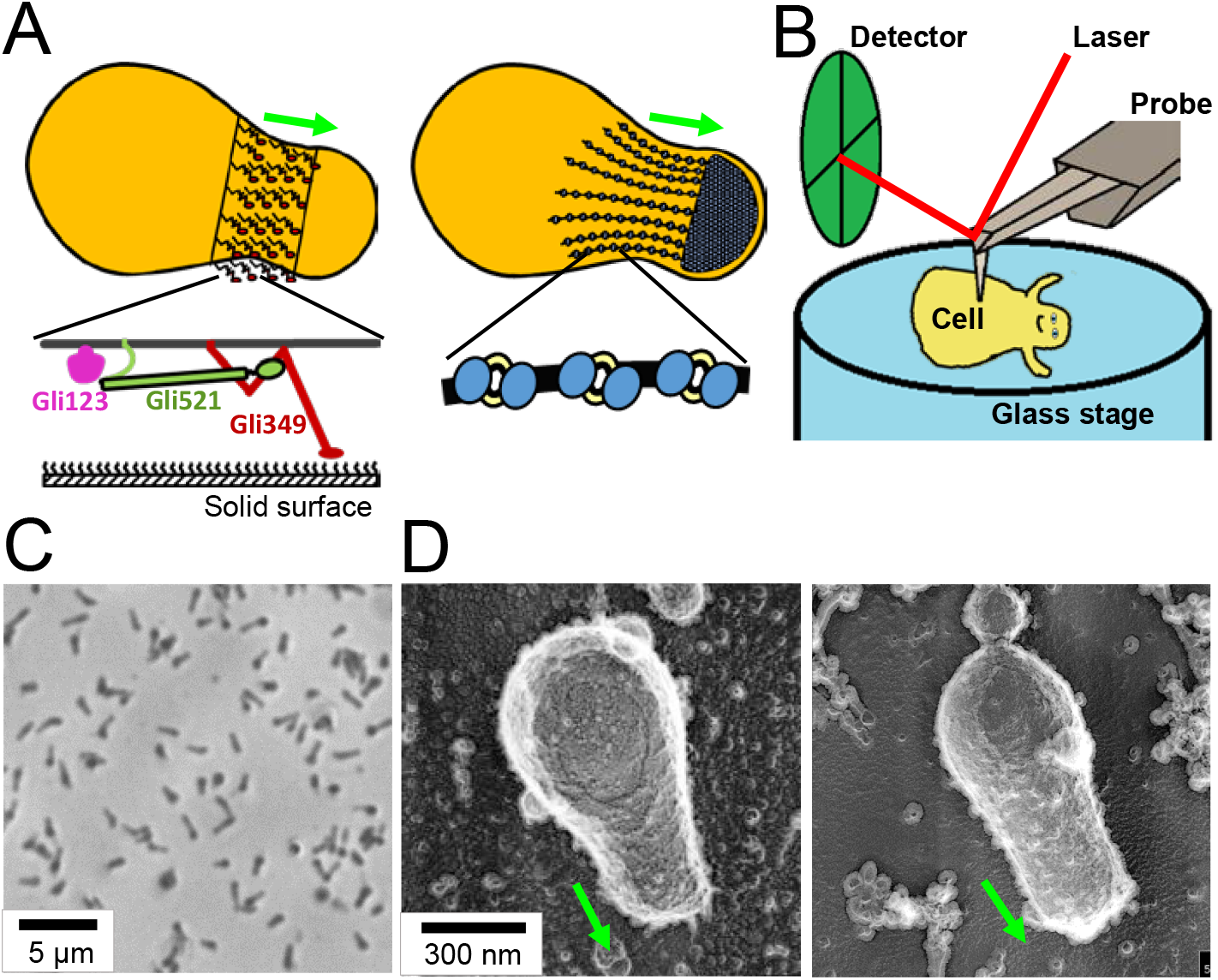
Experimental design and conditions for HS-AFM observation. (A) Schematic illustrations of *M. mobile* gliding machinery. The gliding machinery formed as a protrusion can be divided into surface (left) and internal (right) structures. The surface structure is composed of about 450 units, including three large proteins: Gli123 (purple), Gli521 (green), and Gli349 (red), as shown in the bottom. Gli349 repeatedly catches sialylated oligosaccharides fixed on the solid surface and pulls the cell forward. The internal structure can be divided into a large mass at the cell front and chain structure. The chain structure is composed of particles that have been suggested to evolve from F-type ATPase/synthase. (B) Schematic illustration of *M. mobile* cell being scanned by high-speed atomic force microscopy (HS-AFM). The surface of an immobilized cell on glass substrate (blue) is scanned by an AFM cantilever probe (grey), and the cantilever movement is monitored by a detector (green). (C) Phase-contrast image of *M. mobile* cell on coverslip. Living cells were immobilized onto a coverslip using poly-L-lysine and glutaraldehyde. (D) Quick-freeze, deep-etch EM image of *M. mobile* cells on a cover slip. The cell was immobilized on the coverslip by poly-L-lysine and glutaraldehyde (left) and allowed to glide on the coverslip coated with sialylated oligosaccharides (right). The cell axis and front are indicated by a green arrow (A, D).

High-speed atomic force microscopy (HS-AFM) is a powerful method to monitor the structure and behavior of proteins at the sub-molecular level (29). In this method, a sample placed on a substrate is scanned with a probe and visualized as height information. By performing this process at high speed (~20 frames per second (fps)), the dynamic behavior of samples can be captured while maintaining their active state in aqueous solution. In recent years, this approach has been dramatically improved, and the functional mechanism of more and more proteins has been elucidated *in vitro* through conformational changes (29-33). In addition, HS-AFM has been applied to understand the structures on the cell wall (32) or below the cell membrane (34).

In this study, we succeeded in visualizing the internal structure of *M. mobile* gliding machinery by scanning the surface of cells immobilized on a glass substrate using HS-AFM. The particle structure, a component of the internal structure, showed movements mainly in the right and inward directions relative to the gliding direction of an *M. mobile* cell.

## Results

### Immobilization of living cells on the glass surface

We attempted to visualize the gliding machinery by scanning the upper side of living cells immobilized on the substrate surface (Fig. 1B), since the gliding machinery is arranged around the base of the protruded region (Fig. 1A). Cell suspension in a buffer was placed on a glass substrate reactivated for amino groups and kept for 10 min at 25-28°C. Phase-contrast microscopy showed that the cells adhered to the glass substrate at a density of one cell per approximately 6 μm^2^ (Fig. 1C). When the buffer was replaced by growth medium containing sialylated oligosaccharides (scaffolds for gliding), half of the cells recovered to glide, suggesting that the cells were alive on the glass. Serum included in the medium contained sialylated oligosaccharides conjugated to fetuin, a serum protein. Fetuin was likely adsorbed onto the glass and worked as a scaffold for mycoplasma gliding (22-24, 35).

To observe the shape of immobilized cells, we adopted quick-freeze, deep-etch electron microscopy that visualizes cells under aqueous conditions with nanometer spatial resolution (36, 37). The morphology of immobilized cells (Fig. 1D, left) was not significantly different from that of the gliding cell visualized without any chemical fixation (Fig. 1D right).

### Visualization of immobilized cells by HS-AFM

Next, the cells immobilized on the glass surface were scanned by HS-AFM (Movie S1 and Fig. 2A). A typical *M. mobile* cell with a flask shape was found at a density of a single cell per approximately 100 μm^2^. As can be seen by comparing cell appearance in optical and electron microscopy, the cell images obtained here suggest that cells are characterized by rigidity in the front region (Fig. 1C, D), consistent with previous observations showing an internal rigid “bell” structure (4, 8). The average size of a cell was 0.93 ± 0.33 μm in length and 0.33 ± 0.08 μm in width (n = 20, Fig. 2A). We also measured the height along the long axis of the cell. Two peaks were found: one was near the front end and the other was near the tail end of the cell, consistent with previously reported characteristics of *M. mobile* cells (38, 39).

To visualize the gliding machinery, the cell surface was scanned by HS-AFM at a scanning rate of 300 ms per frame in an area of 300 nm^2^. Interestingly, we found particle structures aligned mostly along the cell axis at the front side of the cells (Fig. 2B). The particle structures appeared when the average tapping force exceeded ~40 pN (see Method). They were aligned at an angle of approximately 4.6° relative to the cell axis (Fig. 2C and D, n = 99 chains from 20 cells). The particle height was approximately 2 nm (Fig. 2E), and the pitches were distributed as 31.5 ± 4.9 nm (Fig. 2F, n = 98) in good agreement with a previous number, 31 nm, measured by electron cryotomography (Fig. 2G)(4). To measure the dimensions of the particles in detail, we collected 19 particle images and averaged them (Fig. 2H). The averaged image showed an elliptical structure, 27.2 nm long and 14.2 nm wide, with two height peaks. The distance between the two peaks of a particle was 10.0 nm. These features were consistent with the results from electron cryotomography (Fig. 2I) (4), showing that the particle structure observed in HS-AFM is identical to the internal structure observed by electron cryotomography.

**FIG 2.**
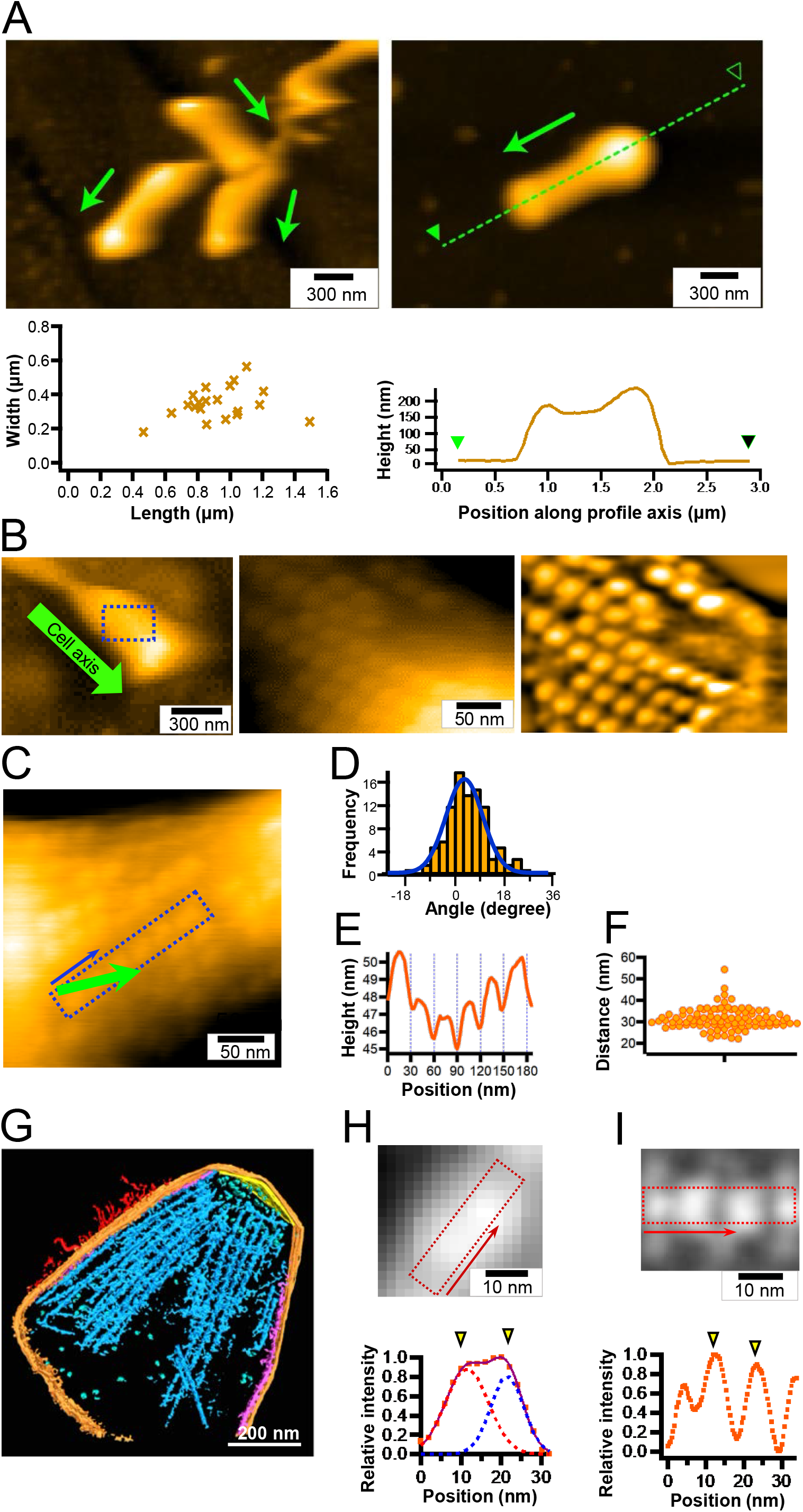
Chain imaging by HS-AFM. (A) Left: Cluster of cells immobilized to glass surface (upper) and distribution of cell dimensions (n = 21)(lower). Right: Height profile along the broken line (upper) is plotted along the green arrow (lower). Cell axis and front are shown by an arrow. (B) Detailed structure of a cell. Left: Whole cell image. The cell axis and front are indicated by a green arrow. Middle: Magnified image of the boxed area of the left panel. Right: The middle panel image was processed with a bandpass filter. (C-F) Image analyses of particles. (C) Cell image featuring a representative chain structure. The cell axis and front are indicated by a green arrow. (D) Distribution of chain angle relative to cell axis fitted by a Gaussian curve (n = 99 chains from 20 cells). (E) Image profile of the boxed area along the direction of blue arrow in panel C. (F) Scatter dot plot for distances between peak positions of chain profile. The average was 31.5 ± 4.9 nm (n = 98). (G) Three-dimensional rendered image for 146-nm-thick slice of permeabilized cell reconstructed by electron cryotomography (4). The surface filamentous structures, cell membrane, undercoating at the front and side membranes, and internal chain are colored red, orange, yellow, and purple, respectively. (H) Averaged image of 19 particle structures from HS-AFM (upper) and image profile of boxed area (lower). The profile (orange squares) was fitted by the sum (purple line) of two Gaussian curves (red and blue). Yellow triangles show peaks of the Gaussian curves. (I) Averaged images of chain structure (blue part in panel G) from electron cryotomography (upper)(4) and image profile of boxed area along the chain axis (lower). Yellow triangles show peaks of Gaussian curves. In all high-speed atomic force microscopy imaging, the surface was scanned left to right for line and lower to upper for image.

### The internal structure of *M. mobile* is detected by HS-AFM from the surface

An *M. mobile* cell has huge proteins on its surface (Fig. 1A, left). To confirm that the particle structures visualized with HS-AFM are not the surface structures, the cell surface was treated with proteinase K, a serine protease with broad specificity, and scanned by HS-AFM. First, we confirmed that *M. mobile* cells gliding on the glass surface were stopped 1 min after the addition of 0.2 mg/mL proteinase K (Fig. S1A), suggesting that the surface proteins involved in the gliding machinery are sensitive to proteinase K. Then, we observed the cell surface by HS-AFM after the immobilized cells were treated with proteinase K for 20 min. The particle structures were observed on the surface of the cell even after proteinase K treatment. The particle pitches of cells with and without proteinase K treatment were 31.2 ± 3.2 (n = 31) and 28.9 ± 3.6 nm (n = 33), respectively (Fig. S1B), showing a significant difference between them (*p* = 0.00651 by Student’s *t*-test). Based on these observations, we concluded that the particle structure detected by HS-AFM was inside the structure, but influenced by the surface treatment with proteinase K, consistent with a previous observation (8).

During the observation of intact cells immobilized on glass surfaces, we observed the removal of the cell membrane by chance, resulting in the exposure of the inside structure. The exposed inside structure showed features similar to the internal jellyfish-like structure of *M. mobile* (4, 8) (Movie S2, Fig. S1C). We compared the features of particle structures before and after the removal of the cell membrane (Fig. S1D). After removal, the height of the particle relative to the background increased, resulting in a clearer appearance than before removal. The particle pitches were 30.3 ± 4.1 and 31.8 ± 7.3 nm before and after removal, respectively, without a statistically significant difference (*p* = 0.277 by Student’s *t*-test). The average heights of particles observed before and after removal of the cell membrane were 257 and 56 nm, respectively, from the lowest position of the image. The difference between them was 201 nm, comparable to the height of *M. mobile* cells (Fig. 2A and Fig. S1D). Therefore, the particles detected before and after cell membrane removal were proposed to be the structure beneath the upper cell membrane and the one on the lower cell membrane facing the glass substrate, respectively. As we could not remove the cell membrane intentionally, we focused on analyzing the internal structure beneath the upper cell membrane.

### Behavior of particle structure detected by HS-AFM

The surface of protrusion of *M. mobile* cell was scanned with a scanning rate of 200 or 330 ms per frame with a scan area of 200 × 200 nm^2^. Projected images were processed using a bandpass filter to improve the image contrast by drift correction and by averaging three sequential images for better signal/noise ratio (Movie S4). In most cases, the particles were difficult to trace over time because of image discontinuity, even when particle images were clear. This is probably due to the stability of the cell immobilized onto the glass surface and damage to the scanning probe. However, we succeeded in tracing the behaviors of individual particles in some videos and used them for further analyses.

### Sodium azide suppressed particle movement

To discuss the behaviors of internal particles, we needed to confirm that the particle movements are caused by ATP hydrolysis on the internal structure. In a previous study, the ATPase activity of the internal structure of *M. mobile* was inhibited by sodium azide (4). The binding activity and gliding speed of “gliding heads”, the gliding machinery isolated from the cell protrusion, were also inhibited by sodium azide (4). In the present study, we examined the effect of sodium azide on the gliding speed of intact *M. mobile* cells. The averaged gliding speed of intact *M. mobile* cells was decreased from 0.77 ± 0.17 to 0.04 ± 0.02 μm/s by the addition of 15.4 mM sodium azide (Fig. 3A, B), suggesting that sodium azide affected the ATPase activity of the internal structure and the force generation for gliding.

**FIG 3.**
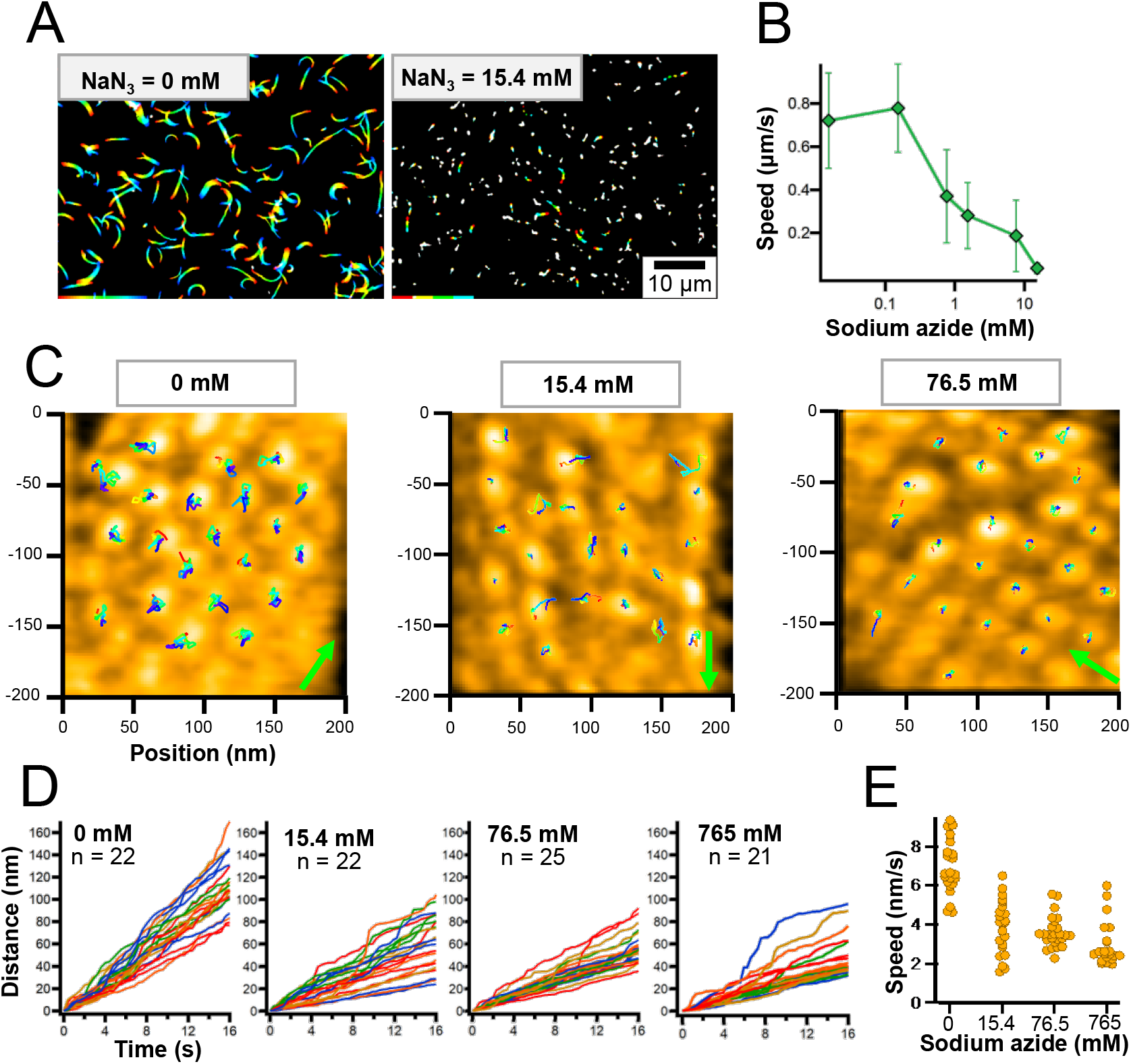
Effects of sodium azide on particle displacements. (A) Rainbow traces of gliding cells for 5 s with and without sodium azide from phase-contrast microscopy. Video frames were overlaid with different colors from red to blue. (B) Gliding speed under various concentrations of sodium azide. Speeds of 2.5-20 s were averaged for 140-223 cells. (C) HS-AFM images with continuous traces of individual particles for 13.2 s. HS-AFM images were processed by bandpass filter, drift correction, and sequential averaging. Particles were traced every 200 ms for no sodium azide, and 330 ms in the presence of sodium azide, as presented by the color change from red to blue. The cell axis and front are indicated by a green arrow. The surface was scanned left to right for line and lower to upper for imaging. Movies are shown as supplemental data as Movies S4, 5, 6, and 7 for imaging in 0, 15.4, 76.5, 765 mM sodium azide. (D) Time course of accumulated moving distances of individual particles under various concentrations of sodium azide. (E) Scatter dot plot of particle speed under various concentrations of sodium azide. Speeds were estimated from a linear fitting of accumulated moving distance.

We then scanned the cell surfaces by HS-AFM in the presence and absence of sodium azide (Movie S4-7). The tracking of the mass center every 200 ms (no azide) or 330 ms (with azide) for 16.2 s showed that most particles were moving independently (Fig. 3C). These movements were significantly reduced by the addition of sodium azide. We calculated the accumulated moving distances and estimated the speeds for the particle movements from a linear fitting of the accumulated moving distance (Fig. 3D, E). At concentrations of 0, 15.4, 76.5, 765 mM sodium azide, the speeds calculated from accumulated moving distances were 6.9 ± 1.4, 3.9 ± 1.4, 3.6 ± 0.8, and 3.0 ± 1.1 nm/s, respectively, suggesting that the movement of particle structures is linked to ATP hydrolysis. Interestingly, in 15.4 mM sodium azide, the particles can be classified as either active or static, and the different types tend to form an adjacent pair in chains (Fig. 3C).

### Particle displacements traced as an image profile

Not all particles moved in the same direction at the same time (Fig. 3C-E), and this feature was more obvious in 15.4 mM sodium azide (Movie S5, Fig. 4A), indicating that the movements were linked to ATP hydrolysis, not caused by artificial drift in the measurements. The addition of sodium azide may allow easier detection of individual movements by reducing some of the movements. Analysis of 27 particles in a 200 × 200 nm^2^ field in the presence of 15.4 mM sodium azide for 23.1 s showed that 19 particles moved distances longer than 6 nm, distinct from other movements. The frequency of such long movements in the whole field was 1.17 events/s (Fig. 4A). Next, we focused on particle movements. Since the particles appeared to move mainly perpendicular to the particle chain in the cell surface plane, the height profile of a box perpendicular to the particle chain was traced over time (Fig. 4B). Six particles did not move (static particle), while the active particles showed remarkable movements, and a returning path for some particles was observed. As shown in “a” panel of Fig. 4B and C, the movements of the particles showed tendency moving 9.1 ± 2.5 nm (n = 15) in the left direction perpendicular to the chain axis and 2.3 ± 3.0 nm (n = 8) on the cytoplasmic side in the Z direction. The profile continued to change for approximately five frames of 330 ms. However, the movement was likely completed in a single 330-ms frame, because the image was profiled after averaging three consecutive video images every 330 ms to reduce image noise. Eleven particles showed returning movements in the video, with similar speeds to their advancing movements, as shown in panels marked “r” in Fig. 4B and C.

**FIG 4.**
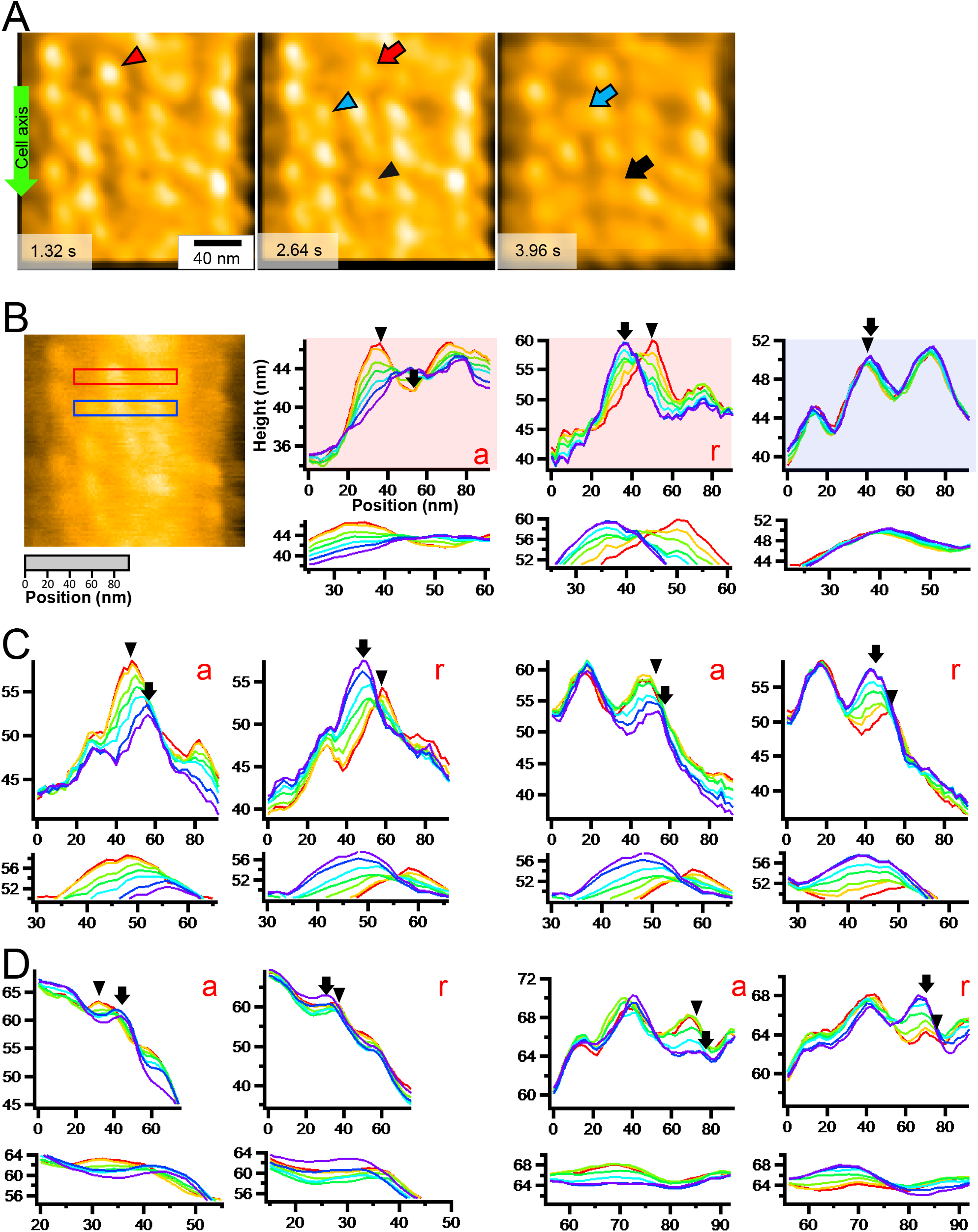
Movements of individual particles. (A) Video frames of particle chains under 15.4 mM sodium azide (Movies S5). The green arrow on the left shows the cell axis and front. The particles with remarkable movements are marked before and after the movements by differently colored triangles and arrows, respectively. Particles moved to the left relative to the gliding direction. (B) Consecutive image profile of active and static particles. Left image: Raw image of video frame showing areas profiled for active (red) and static (blue) particles. Right graphs: Image profiles of active (red background) and static (blue background) particles every 330 ms for 1.98 s. (C) Consecutive image profiles showing particle movements every 330 ms for 1.98 s in 15.4 mM sodium azide. (D) Consecutive image profiles showing particle movements every 200 ms for 1.2 s without sodium azide (Movies S4). (B, C, D) Consecutive profiles of each frame from red to purple. Advancing (a) and returning (r) movements are presented. Peak positions of focusing particles are marked by a triangle and an arrow, respectively, for the initial and the end time points. Distances between peaks before and after movement are marked by a triangle and an arrow, respectively; these were manually measured for statistical analysis of particle movements. The profile of heights and positions is presented with a common X-Y-scale in the lower panel for each data set.

Next, particle movements perpendicular to the cell axis were searched in the absence of sodium azide. Observation of 21 particles for 16.6 s showed that movements longer than 6 nm appeared at a frequency of 2.17 events/s (Movie S4 and Fig. 4D). The distance moved was 8.0 ± 1.9 nm (n = 24) in the left direction perpendicular to the axis of the chain alignment within 200 ms and 2.0 ± 1.9 nm (n = 18) on the cytoplasmic side in the Z direction (Fig. 4D).

### Particle displacements traced as a positional distribution

To study the direction of movements of the particles on the membrane surface statistically, the distributions of the particles as the mass center were analyzed every 200 and 330 ms for observations in the absence and presence of sodium azide, respectively (Fig. 5A and Movies S5-7). The faster-scan speed for the observation in the absence of sodium azide was applied, as we assumed that the particles moved faster in this condition. However, this difference in the scanning speed should not affect the conclusion, because no difference was found, even when the analysis was performed using 400 ms intervals for the measurements without sodium azide (Fig. S2). Analysis showed that the distributions were larger in the presence of 15.4 mM and smaller at 76.5 and 765 mM than in the absence of sodium azide (Fig. 5A). Next, we measured the distributions of three distances (Fig. 5B): the particle position to the chain axis (Fig. 5C), the distance to the adjacent particle (Fig. 5D), and the distance to the adjacent particle projected to the chain axis (Fig. 5E). These results are schematically summarized (Fig. 5B), suggesting that movements perpendicular to the chain axis of the particles (presented as movement “c” in Fig. 5) should be present but not easy to detect in the absence of sodium azide; they were observed more clearly when the frequency of movements was reduced by sodium azide, and they were inhibited under high concentrations of sodium azide.

**FIG 5.**
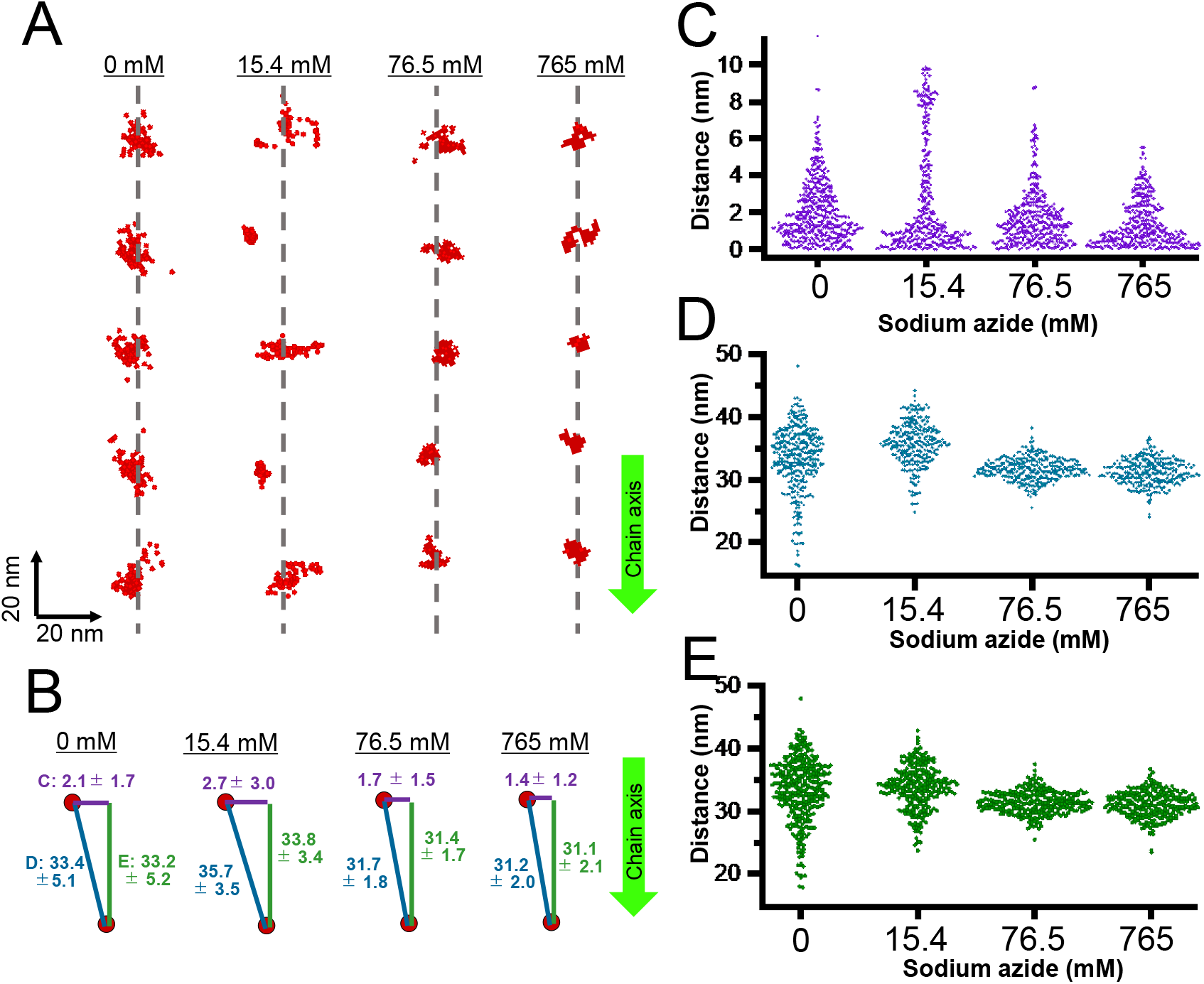
Analyses of particle distribution. (A) Distribution of particles in chain. The particle positions and the axis of the particle positions are indicated by red dots and grey dashed lines, respectively. The particle positions were detected every 200 and 330 ms, respectively, without and with sodium azide at 82, 66, 70, and 66 points under 0, 15.4, 76.5, and 765 mM sodium azide, respectively. The axis of particle positions was determined by a linear approximation of the average position of each particle. (B) Schematic illustration of three distances with average and standard deviation (SD) values in nm. The particle position to the chain axis (C, purple), the distance to the adjacent particle (D, blue), and the distance to the adjacent particle projected to the chain axis (E, green) are shown. Bar lengths are not to scale. Movies S4-7 were analyzed. The chain axis is indicated by a green arrow pointing mostly to the cell front in panels (A) and (B).

## Discussion

### Internal structure was traced from the outside surface

The particle features traced by HS-AFM in this study were consistent with those of the internal structure reported in previous studies (Fig. 2) (4, 8), suggesting that HS-AFM visualized the internal structure. The large surface proteins Gli521, Gli349, and Gli123 exist on the cell surface of *M. mobile* as components of the gliding machinery (10, 13-16, 18, 20, 39, 40). A group of surface proteins, Mvsps, which are responsible for antigenic variations, also exist on the cell surface (41,42). These surface proteins may interfere with probing the internal structure from the surface. However, the chain structures observed by HS-AFM did not show significant differences before and after protease treatment of the cells (Fig. S1 B). Furthermore, similar structures were observed before and after mechanical removal of the cell membrane (Fig. S1C, D). These results showed that the particles traced by HS-AFM were not on the surface structure, but inside the cell. The surface structure, composed of mainly large filamentous proteins, may be too thin and/or mobile to be detected by the current scanning performance of HS-AFM on the cell membrane (12-14, 18). The lack of a peptidoglycan layer should be advantageous for visualizing the inside structure, due to the lack of stiffness (36-38). Moreover, the internal structure should be sufficiently stiff and positioned beneath the cell membrane, reminiscent of cortical actin in animal cells (34).

### Effects of sodium azide

Sodium azide inhibits many ATPases by blocking ADP release (43). In *M. mobile* gliding, the reagent inhibited cell gliding (Fig. 3A, B) and the isolated gliding machinery (4). Particle behaviors became more visible in the presence of 15.4 mM sodium azide. Under this condition, cell gliding was reduced to 20 times slower than the original, suggesting that ATP hydrolysis occurred 20 times less frequently. If the particles move in a rapid and independent manner, it may be difficult to trace the movements of individual particles. However, if the reaction was partially inhibited by 15.4 mM sodium azide, most particles may be in their home position, while some particles move to another position. In this case, the movements could be traced easily. This assumption is supported by the observation that the particle distances between neighboring particles are 1.7–2.5 nm shorter under high concentrations of sodium azide than those without the reagent (Fig. 5D). A previous study based on electron microscopy showed that the particle distances in the ADP and unbound forms were approximately 2 nm shorter than those in the AMPPNP, ADP-V_i_, and ADP-AlF_x_ states (4). As sodium azide is thought to inhibit the release of ADP (43), the changes in particle distance observed in the present study are consistent with the results of electron microscopy (Fig. 5B)(4).

### Particle behavior in the gliding mechanism

The particles moved approximately 9 nm to the right of the gliding direction and 2 nm to the cytoplasmic side within 330 ms (Fig. 6). This movement may be coupled with the transition from ADP or unbound form to ATP or the ADP/P_i_ form (4). Considering the fact that the particles are structurally linked to the surface structures of the gliding machinery (4), the movements observed in the present study are likely involved in the gliding mechanism.

**FIG 6.**
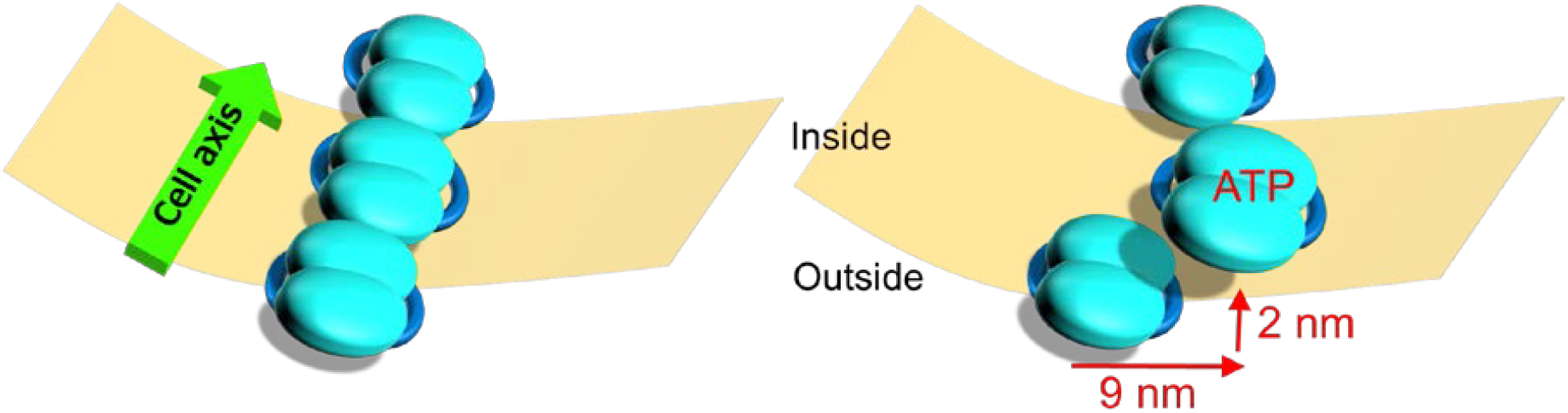
Schematic illustration of particle movement in *M. mobile* visualized by HS-AFM. The internal chain of the gliding machinery and cell membrane are indicated by blue objects and a beige plate, respectively. Here, we focus on the particle chain lining the lower side of cell membrane, while we scanned mostly the particle chain beneath the upper side of cell membrane in this study. The left and right panels show the particles before and after the advancing movement, respectively. The central particle moves as an ATP-or ADP/Pi-bound form, to the right and inner sides for a distance of 9 and 2 nm, respectively.

Previous studies have reported that the step size of *M. mobile* is approximately 70 nm under no load and adjustable to various loads (17, 25, 28, 44). The surface structure contains two large proteins with dimensions comparable to the step size, that is, the Gli349 “leg” that catches the scaffold and the Gli521 “crank” that transmits force for gliding are 100 and 120 nm long, respectively (Fig. 1A) (12, 13, 18). In the present study, we could not detect conformational changes in the internal structure with length comparable to the step size. Therefore, the movements occurring in the internal structure should be amplified through the huge protein molecules on the surface or through an unknown structure that connects the internal and surface structures (Fig. 1A, 2G). This assumption can explain the previous observation that the single leg exerts a force of 1.5 pN, a few times smaller than motor proteins (17), assuming elastic components are equipped in the huge surface complex.

In a previous study, *M. mobile* gliding showed a leftward directional change of about 8.5° with 1-μm cell progress (27). This is consistent with the observation that the particle movements are pointed to the right, relative to the gliding direction (Fig. 6). Otherwise, the tilting of the chain axis about 4.6° from the cell axis may cause a directional change in gliding (Fig. 2D).

To elucidate the mechanism of *M. mobile* gliding, we need to further visualize the behaviors and structures of the machinery in detail, including those of both internal and surface structures. The combination of electron microscopy and HS-AFM may provide better insights in the near future.

## Materials and Methods

### Cell preparation

A mutant strain (*gli521*[P476R]) of *M. mobile* 163K (ATCC43663) activated for binding (17, 19, 45) was grown in Aluotto medium at 25– 28°C, as previously described (36, 39). Cultured cells were collected by centrifugation at 12,000 × *g* for 4 min at 25–28°C and suspended in phosphate-buffered saline with glucose (PBS/G) consisting of 75 mM sodium phosphate (pH 7.3), 68 mM NaCl, and 10 mM glucose (17, 22, 26, 27). This process was repeated twice, and finally the cells were resuspended in PBS/G to a 20-fold density of the original culture.

### Gliding analyses

A tunnel chamber assembled as previously described (3-mm interior width, 22-mm length, 40-μm wall thickness) was treated with Aluotto medium for 15 min at 25–28°C (17, 26), and then the medium was replaced by PBS/G. The cell suspension was inserted into the tunnel chamber with video recording. PBS/G was replaced with PBS/G containing 0.2 mg/mL proteinase K (Qiagen N. V., Hilden, Germany) or various concentrations of sodium azide, as necessary.

### Cell immobilization on the glass surface

A glass slide was treated with saturated KOH-ethanol solution for 15 min and washed 10 times with water. For analyses with an imaging rate of 1000 and 330 ms per frame, the glass was treated with 0.1% poly-L-lysine for 5 min. After the solution was removed, the glass was washed with water and dried. Then, the glass was treated with 0.1% glutaraldehyde for 5 min, washed with water, and covered with PBS/G. For analyses with an imaging rate of 200 ms per frame, the glass was treated with sandpaper, saturated with KOH-ethanol solution for 15 min, washed 10 times with water, and then dried. The washed glass was treated with 1000-fold diluted 3-aminopropyldiethoxymethylsilane for 5 min at 25–28°C, washed, and treated with glutaraldehyde as described above. Finally, the cell suspension was placed onto the glass substrate and left for 10 min at 25–28°C.

### Microscopy

To examine the immobilizing conditions using phase-contrast microscopy, the glass slide was assembled into a tunnel chamber (15). The cell suspension was loaded into the tunnel, kept for 10 min at 25–28°C, washed with PBS/G, and observed by phase-contrast microscopy IX71 (Olympus, Tokyo, Japan)(17, 23, 27). To analyze the immobilizing conditions, quick-freeze deep-etch electron microscopy, fixation, and washing were performed on the coverslip. When the cells were frozen without immobilization, we followed the procedure for the electron microscopy method described previously (36, 37). Briefly, the cells on the glass were pressed against a copper block cooled with liquid helium and frozen. Then, the frozen sample was fractured and etched to expose it. Subsequently, the exposed surface was shadowed with platinum to create a replica membrane, which was observed under a JEM-1010 transmission electron microscope (JEOL, Tokyo, Japan) at 80 kV, equipped with a FastScan-F214 (T) charge-coupled device (CCD) camera (TVIPS, Gauting, Germany).

### Observation by HS-AFM

Imaging was performed with a laboratory-built HS-AFM in tapping mode (46, 47). Small cantilevers (BLAC10DS-A2, Olympus), with a resonant frequency of ~0.5 MHz in water, a quality factor (*Q*_c_) of ~1.5 in water, and a spring constant (*k*_c_) of ~0.08 N/m were used. The cantilever’s free oscillation amplitude (*A*_0_) and set-point amplitude (*A*_sp_) were set at ~2.5 nm and ~0.8 × *A*_0_, respectively. From this condition, the average tapping force *<F>* can be approximated as ~40 pN using the following equation: 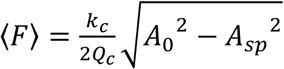 For searching cells, the sample was scanned at an imaging rate of 1000 ms per frame in an area of 3000 × 3000 nm^2^ with 150 × 150 pixels. To observe the particle structure, the cell surface was scanned with an imaging rate of 330 or 200 ms per frame in an area of 200 × 200 nm^2^ with 100 × 100 pixels.

### Video analyses

To trace particles in the XY plane, videos were processed by three methods (Movies S3-7). (i) The image contrast was improved by a bandpass filter. (ii) Image drifts were corrected by a plugin, “align slices in stack” (48), equipped with ImageJ. (iii) Image noises were removed by averaging three consecutive slices. Then, each particle image was cropped, binarized, and traced for the mass center. Here, the threshold for binarization was determined independently for each particle of interest. The cell axes in Fig. 2 were determined by fitting a cell image as an ellipse. All analyses were performed with ImageJ 1.52A. Image averaging of particles was performed using EMAN, version 2.3.

## Acknowledgments

We appreciate Yuya Sasajima at Osaka City University for helpful discussions. This work was supported by Grants-in-Aid for Scientific Research (B) and (A) (MEXT KAKENHI, Grant Numbers JP24390107, JP17H01544), JST CREST (Grant Number JPMJCR19S5), Osaka City University (OCU) Strategic Research Grant 2018 for top priority research, and by a Grant-in-Aid for the Fugaku Trust for Medicinal Research to MM.

---

**FIG S1 Protease treatment and unroofing confirm that particles are intracellular structures.** (A) Rainbow traces of gliding cell for 5 s starting 20 s before and 60 s after the addition of 0.2 mg/ml proteinase K marked from red to purple over time. (B) Processed HS-AFM image of particle structures on a cell surface without (left) and with (middle) proteinase K treatment and scatter dot plot of distances between particles along each chain axis (right). Particle distances on cells with and without proteinase K treatment were 30.1 ± 6.1 nm (n = 35) and 31.2 ± 3.2 nm (n = 31), respectively (Student’s *t*-test: *p* = 0.328). (C) Time course images of HS-AFM scanning of cell membrane removal. The cell membrane started to be broken at 2 s, and the internal structure was completely exposed at 23 s after the cell was focused (t = 0)(Movies S2). Scanning area, 500 × 500 nm^2^ with 150 × 150 pixels; frame rate, 1000 ms per frame. (D) Magnified HS-AFM image of particle structures before (left) and after (middle) cell membrane removal and scatter dot plot of distances between particles along each chain axis (right). The color gauge on the right side of each figure shows the scale of the relative height (the height is presented by adjusting the lowest point in panel D until it becomes 0). The averaged height of the observation surface after removal of cell membrane was 201 nm lower than before removal (257 and 56 nm for before and after, respectively). The distances between neighboring particles before and after cell membrane removal were 30.3 ± 4.1 (n = 36) and 31.8 ± 7.3 nm (n = 40), respectively (*p* = 0.277). The cell axis and front are indicated by a green arrow in panels (B-D).

**FIG S2 Particle distribution without sodium azide analyzed using different time intervals.** Distribution of particles in a chain (A), the particle position to the chain axis (B), the distance to the adjacent particle (C), and the distance to the adjacent particle projected to the chain axis (D) were analyzed every 200, 400, and 600 ms.

**Movie S1 HS-AFM movie searching for *M. mobile* cells.** An *M. mobile* cell immobilized to the substrate surface was searched by recording at 1 fps. The video was played at a speed of 2×. The scanning field was 3 × 3 μm^2^ with 100 × 100 pixels. A cell appeared around 6 s and moved around the center of the field at 13 s. The cell front is directed to the lower right.

**Movie S2 HS-AFM movie showing removal of the cell membrane.** The upper membrane of a cell scanned at 3 fps was removed at approximately 20 s. The video was played at 5× speed. The scanning field was 500 × 500 nm^2^ with 150 × 150 pixels. The cell front is directed to the upper right.

**Movie S3 Effects of image processing on HS-AFM movies of the cell surface.** The original movie (upper left) was processed using a bandpass filter (upper right), bandpass filter + drift correction (lower left), and bandpass filter + drift correction + sequential averaging (lower right). The video was played at 3.3× speed. The cell front was directed to the lower portion of the frame.

**Movie S4 HS-AFM movie showing particle movements.** The cell surface was scanned at 5 fps. The scanning field was 200 × 200 nm^2^ with 100 × 100 pixels. The video was played at 2× speed. The cell front is directed to the upper right.

**Movie S5 HS-AFM movie showing particle movements under 15.4 mM sodium azide.** The cell surface was scanned at 3 fps. The scanning field was 200 × 200 nm^2^ with 100 × 100 pixels. The video was played at 3.3× speed. The cell front was directed to the lower portion of the frame.

**Movie S6 HS-AFM movie showing particle movements under 76.5 mM sodium azide.** The cell surface was scanned at 3 fps. The scanning field was 200 × 200 nm^2^ with 100 × 100 pixels. The video was played at 3.3× speed. The cell front is directed to the upper left.

**Movie S7 HS-AFM movie showing particle movements under 765 mM sodium azide.** The cell surface was scanned at 3 fps. The scanning field was 200 × 200 nm^2^ with 100 × 100 pixels. The video was played at 3.3× speed. The cell front is directed to the upper left.

## References

1. Miyata M, Robinson RC, Uyeda TQP, Fukumori Y, Fukushima SI, Haruta S, Homma M, Inaba K, Ito M, Kaito C, Kato K, Kenri T, Kinosita Y, Kojima S, Minamino T, Mori H, Nakamura S, Nakane D, Nakayama K, Nishiyama M, Shibata S, Shimabukuro K, Tamakoshi M, Taoka A, Tashiro Y, Tulum I, Wada H, Wakabayashi KI. 2020. Tree of motility - A proposed history of motility systems in the tree of life. Genes Cells 25:6–21.

2. Nakamura S, Minamino T. 2019. Flagella-driven motility of bacteria. Biomolecules 9: 31337100.

3. Tulum I, Kimura K, Miyata M. 2020. Identification and sequence analyses of the gliding machinery proteins from *Mycoplasma mobile*. Sci Rep 10:3792.

4. Nishikawa M, Nakane D, Toyonaga T, Kawamoto A, Kato T, Namba K, Miyata M. 2019. Refined mechanism of *Mycoplasma mobile* gliding based on structure, ATPase activity, and sialic acid binding of machinery. mBio 10:e02846–02819.

5. Hamaguchi T, Kawakami M, Furukawa H, Miyata M. 2019. Identification of novel protein domain for sialyloligosaccharide binding essential to *Mycoplasma mobile* gliding. FEMS Microbiol Lett 366:fnz016.

6. Miyata M, Hamaguchi T. 2016. Prospects for the gliding mechanism of *Mycoplasma mobile*. Curr Opin Microbiol 29:15–21.

7. Tulum I, Yabe M, Uenoyama A, Miyata M. 2014. Localization of P42 and F_1_-ATPase alpha-subunit homolog of the gliding machinery in *Mycoplasma mobile* revealed by newly developed gene manipulation and fluorescent protein tagging. J Bacteriol 196:1815–1824.

8. Nakane D, Miyata M. 2007. Cytoskeletal “jellyfish” structure of *Mycoplasma mobile*. Proc Natl Acad Sci U S A 104:19518–19523.

9. Miyata M. 2010. Unique centipede mechanism of *Mycoplasma* gliding. Annu Rev Microbiol 64:519–537.

10. Uenoyama A, Miyata M. 2005. Identification of a 123-kilodalton protein (Gli123) involved in machinery for gliding motility of *Mycoplasma mobile*. J Bacteriol 187:5578–5584.

11. Beven L, Charenton C, Dautant A, Bouyssou G, Labroussaa F, Skollermo A, Persson A, Blanchard A, Sirand-Pugnet P. 2012. Specific evolution of F1-like ATPases in mycoplasmas. PLoS One 7:e38793.

12. Lesoil C, Nonaka T, Sekiguchi H, Osada T, Miyata M, Afrin R, Ikai A. 2010. Molecular shape and binding force of *Mycoplasma mobile’s* leg protein Gli349 revealed by an AFM study. Biochem Biophys Res Commun 391:1312–1317.

13. Adan-Kubo J, Uenoyama A, Arata T, Miyata M. 2006. Morphology of isolated Gli349, a leg protein responsible for *Mycoplasma mobile* gliding via glass binding, revealed by rotary shadowing electron microscopy. J Bacteriol 188:2821–2828.

14. Metsugi S, Uenoyama A, Adan-Kubo J, Miyata M, Yura K, Kono H, Go N. 2005. Sequence analysis of the gliding protein Gli349 in *Mycoplasma mobile*. Biophysics (Nagoya-shi) 1:33–43.

15. Uenoyama A, Kusumoto A, Miyata M. 2004. Identification of a 349-kilodalton protein (Gli349) responsible for cytadherence and glass binding during gliding of *Mycoplasma mobile*. J Bacteriol 186:1537–1545.

16. Kusumoto A, Seto S, Jaffe JD, Miyata M. 2004. Cell surface differentiation of *Mycoplasma mobile* visualized by surface protein localization. Microbiology 150:4001–4008.

17. Mizutani M, Tulum I, Kinosita Y, Nishizaka T, Miyata M. 2018. Detailed analyses of stall force generation in *Mycoplasma mobile* gliding. Biophys J 114:1411–1419.

18. Nonaka T, Adan-Kubo J, Miyata M. 2010. Triskelion structure of the Gli521 protein, involved in the gliding mechanism of *Mycoplasma mobile*. J Bacteriol 192:636–642.

19. Uenoyama A, Seto S, Nakane D, Miyata M. 2009. Regions on Gli349 and Gli521 protein molecules directly involved in movements of *Mycoplasma mobile* gliding machinery, suggested by use of inhibitory antibodies and mutants. J Bacteriol 191:1982–1985.

20. Seto S, Uenoyama A, Miyata M. 2005. Identification of a 521-kilodalton protein (Gli521) involved in force generation or force transmission for *Mycoplasma mobile* gliding. J Bacteriol 187:3502–3510.

21. Chen J, Neu J, Miyata M, Oster G. 2009. Motor-substrate interactions in *Mycoplasma* motility explains non-Arrhenius temperature dependence. Biophys J 97:2930–2938.

22. Kasai T, Hamaguchi T, Miyata M. 2015. Gliding motility of *Mycoplasma mobile* on uniform oligosaccharides. J Bacteriol 197:2952–2957.

23. Kasai T, Nakane D, Ishida H, Ando H, Kiso M, Miyata M. 2013. Role of binding in *Mycoplasma mobile* and *Mycoplasma pneumoniae* gliding analyzed through inhibition by synthesized sialylated compounds. J Bacteriol 195:429–435.

24. Nagai R, Miyata M. 2006. Gliding motility of *Mycoplasma mobile* can occur by repeated binding to *N*-acetylneuraminyllactose (sialyllactose) fixed on solid surfaces. J Bacteriol 188:6469–6475.

25. Kinosita Y, Miyata M, Nishizaka T. 2018. Linear motor driven-rotary motion of a membrane-permeabilized ghost in *Mycoplasma mobile*. Sci Rep 8:11513.

26. Tanaka A, Nakane D, Mizutani M, Nishizaka T, Miyata M. 2016. Directed binding of gliding bacterium, *Mycoplasma mobile,* shown by detachment force and bond lifetime. mBio 7:00455–00416.

27. Morio H, Kasai T, Miyata M. 2016. Gliding direction of *Mycoplasma mobile*. J Bacteriol 198:283–290.

28. Kinosita Y, Nakane D, Sugawa M, Masaike T, Mizutani K, Miyata M, Nishizaka T. 2014. Unitary step of gliding machinery in *Mycoplasma mobile*. Proc Natl Acad Sci USA 111:8601–8606.

29. Ando T. 2018. High-speed atomic force microscopy and its future prospects. Biophys Rev 10:285–292.

30. Kodera N, Noshiro D, Dora SK, Mori T, Habchi J, Blocquel D, Gruet A, Dosnon M, Salladini E, Bignon C, Fujioka Y, Oda T, Noda NN, Sato M, Lotti M, Mizuguchi M, Longhi S, Ando T. 2020. Structural and dynamics analysis of intrinsically disordered proteins by high-speed atomic force microscopy. Nat Nanotechnol doi:10.1038/s41565-020-00798-9.

31. Kodera N, Ando T. 2020. High-speed atomic force microscopy to study myosin motility. Adv Exp Med Biol 1239:127–152.

32. Yamashita H, Taoka A, Uchihashi T, Asano T, Ando T, Fukumori Y. 2012. Single-molecule imaging on living bacterial cell surface by high-speed AFM. J Mol Biol 422:300–309.

33. Kodera N, Yamamoto D, Ishikawa R, Ando T. 2010. Video imaging of walking myosin V by high-speed atomic force microscopy. Nature 468:72–76.

34. Zhang Y, Yoshida A, Sakai N, Uekusa Y, Kumeta M, Yoshimura SH. 2017. In vivo dynamics of the cortical actin network revealed by fast-scanning atomic force microscopy. Microscopy (Oxf) 66:272–282.

35. Jaffe JD, Miyata M, Berg HC. 2004. Energetics of gliding motility in *Mycoplasma mobile*. J Bacteriol 186:4254–4261.

36. Tulum I, Tahara Y, Miyata M. 2019. Peptidoglycan layer and disruption processes in *Bacillus subtilis* cells visualized using quick-freeze, deep-etch electron microscopy. Microscopy (Oxf) 68:441–449.

37. Miyata M, Petersen JD. 2004. Spike structure at the interface between gliding *Mycoplasma mobile* cells and glass surfaces visualized by rapid-freeze-and-fracture electron microscopy. J Bacteriol 186:4382–4386.

38. Nakane D, Miyata M. 2012. *Mycoplasma mobile* cells elongated by detergent and their pivoting movements in gliding. J Bacteriol 194:122–130.

39. Miyata M, Yamamoto H, Shimizu T, Uenoyama A, Citti C, Rosengarten R. 2000. Gliding mutants of *Mycoplasma mobile:* relationships between motility and cell morphology, cell adhesion and microcolony formation. Microbiology 146:1311–1320.

40. Wu HN, Miyata M. 2012. Whole surface image of *Mycoplasma mobile,* suggested by protein identification and immunofluorescence microscopy. J Bacteriol 194:5848–5855.

41. Wu HN, Kawaguchi C, Nakane D, Miyata M. 2012. “Mycoplasmal antigen modulation,” a novel surface variation suggested for a lipoprotein specifically localized on *Mycoplasma mobile*. Curr Microbiol 64:433–440.

42. Adan-Kubo J, Yoshii SH, Kono H, Miyata M. 2012. Molecular structure of isolated MvspI, a variable surface protein of the fish pathogen *Mycoplasma mobile*. J Bacteriol 194:3050–3057.

43. Bowler MW, Montgomery MG, Leslie AG, Walker JE. 2006. How azide inhibits ATP hydrolysis by the F-ATPases. Proc Natl Acad Sci U S A 103:8646–8649.

44. Miyata M, Ryu WS, Berg HC. 2002. Force and velocity of *Mycoplasma mobile* gliding. J Bacteriol 184:1827–1831.

45. Uenoyama A, Miyata M. 2005. Gliding ghosts of *Mycoplasma mobile*. Proc Natl Acad Sci USA 102:12754–12758.

46. Uchihashi T, Kodera N, Ando T. 2012. Guide to video recording of structure dynamics and dynamic processes of proteins by high-speed atomic force microscopy. Nat Protoc 7:1193–1206.

47. Ando T, Kodera N, Takai E, Maruyama D, Saito K, Toda A. 2001. A high-speed atomic force microscope for studying biological macromolecules. Proc Natl Acad Sci U S A 98:12468–12472.

48. Tseng Q, Duchemin-Pelletier E, Deshiere A, Balland M, Guillou H, Filhol O, Thery M. 2012. Spatial organization of the extracellular matrix regulates cell-cell junction positioning. Proc Natl Acad Sci U S A 109:1506–1511.

